# VoroTIDE: Protein domain segmentation through independent hierarchical searches using Voronoi tessellation and physical self-containment

**DOI:** 10.64898/2026.10.18.759833

**Authors:** Quoc Khang Le, Guillaume Postic, Eric Angel, Fariza Tahi

## Abstract

Protein domains are commonly regarded as structurally separated, functionally relevant, and evolutionarily separable units. Although classification systems may assign different domain boundaries to the same structure, structural separation remains a shared criterion, and many domain parsers assess it through inter-residue distances, contact numbers, or contact densities. However, the Voronoi contact area, which is widely used to study protein–protein interactions and predict contact sites, has seen limited use in domain parsing. There are two potential challenges to this problem. First, unlike in protein-protein interaction, partitions are not known in advance and must be identified before their interface can be evaluated. Second, the contact area depends on fragment size and internal packing, making it difficult to compare candidate partitions. Here, we introduce VoroTIDE (Voronoi-based Two-way Independent Domain Exploration), a domain parser based on all-heavy-atom Voronoi contact areas. By normalizing the contact area for fragment size and internal packing, and incorporating residue hydropathy, we define a pairwise self-containment score F. VoroTIDE applies F in two independent hierarchical searches: top-down recursive splitting until no admissible division remains, and bottom-up merging of elementary structural units until the entire structure becomes a single fragment. Intermediate partitions from both searches are ranked by a global partition quality score G, derived on the same principle. On two CATH–ECOD consensus benchmarks, under balanced evaluation, VoroTIDE achieved the strongest performance among all tested criterion-based parsers and was comparable to Chainsaw, the best-performing deep learning method tested. These results show that the Voronoi-based criterion can yield competitive domain parsing.

**Software availability:** https://github.com/LeKhang97/VoroTIDE

**Statement of importance:** Protein domains are functional units that are structurally separated. Therefore, one natural solution is to identify domains by measuring the Voronoi contact areas among potential fragments. However, potential fragments are not known in advance, and distinguishing candidate partitions is also challenging. This study shows that VoroTIDE addresses both problems using Voronoi-based criteria, providing competitive domain-parsing performance and demonstrating that such criteria can support both boundary discovery and partition assessment.

## Introduction

Protein domains are compact structural units with partial independence in folding, evolution, or function (Wetlaufer 1973, Porter and Rose 2012). Therefore, identifying protein domains is important for structural classification, evolutionary and functional analyses, and structure-based design (Sillitoe *et al*. 2020, Wang *et al*. 2021, Durham *et al*. 2025, Kandathil *et al*. 2025). As the number of experimentally determined and predicted protein structures continues to grow, many automated methods for domain parsing have been developed and become increasingly important (Lau *et al*. 2024).

Protein domain parsers can be broadly grouped into methods based on explicit structural criteria (criterion-based methods) and deep-learning methods. For criterion-based methods, there are PUU (Holm and Sander 1994), DomainParser (Xu, Xu, and Gabow 2000), PDP (Alexandrov and Shindyalov 2003), SWORD (Postic et al. 2017), DDOMAIN (Zhou, Xue, and Zhou 2007) and UniDoc (Zhu et al. 2023) that detect weak structural coupling from contacts, distances, or interaction potentials. In contrast, deep-learning methods infer domain organization using trained models: Chainsaw (Wells et al. 2024) predicts residue-pair same-domain probabilities, Merizo (Lau, Kandathil, and Jones 2023) predicts domain masks, and DPAM-AI (Durham et al. 2025) integrates learned structural signals with predicted aligned errors and homology information. These methods have achieved strong performance on annotated benchmarks, but their boundary decisions depend on learned representations. Methods based on explicit criteria remain useful when the structural basis of each decision needs to be examined.

Many criterion-based parsers represent contacts using distance cutoffs or distance-based functions. Such representations quantify proximity but do not account for occlusion by atoms between two residues, nor do they directly measure the geometric extent of an interface. Voronoi geometry provides a complementary description in which contact areas are derived from shared boundaries between atomic cells (Poupon 2004, Cazals et al. 2006). Surface-based criteria have a long history in domain parsing: Wodak and Janin (1981) identified candidate boundaries from minima in interfacial surface area (Wodak and Janin 1981), and STRUDL used weighted Voronoi contact areas defined by radical planes (Wernisch, Hunting, and Wodak 1999). DePot used a residue lattice derived from Delaunay tessellation, the geometric dual of a Voronoi tessellation (Taylor and Vaisman 2006). These precedents motivate further investigation of tessellation-based contacts as explicit criteria for protein-domain segmentation.

Nevertheless, Voronoi-based approaches have remained limited in protein-domain parsing due to several challenges. One challenge is that the two problems pose different geometric tasks. In protein–protein interface analysis, the interacting partners are known a priori, so the interface to be characterized is already defined. In domain parsing, by contrast, the partition itself is unknown and candidate fragments must be discovered during the search. Moreover, raw Voronoi interface areas cannot be compared directly across candidate fragments because they depend on fragment size and internal packing. Applying Voronoi geometry to domain parsing therefore requires both a search over candidate partitions and a normalized criterion for comparing them.

STRUDL has addressed these issues. Using a Kernighan–Lin heuristic on a Voronoi contact graph of a structure containing *N* residues, STRUDL searches for bipartitions in which one fragment has a target size of *k* residues, using Voronoi contact area as the objective function. Among bipartitions with the same target size, STRUDL chose the one with the lowest contact area. Among the chosen bipartitions of different target sizes, the program normalizes the minimum contact area to *k*(*N-k)*, then selects the bipartition with the lowest normalized contact area as the candidate. This candidate bipartition is subsequently accepted or rejected based on additional criteria that describe relative inter-fragment contact and internal compactness. Accepted partitions are then processed recursively to obtain a partition with more fragments.

STRUDL demonstrated that the Voronoi contact area can be applied effectively to domain parsing. However, while its candidate generation used the Voronoi contact area, its split acceptance relied on additional criteria: inter-fragment contact and internal compactness. This required additional calibration, thereby increasing the risk of overfitting, while also separating the criterion used to generate candidate splits from that used to accept them. This raises the question of whether the same Voronoi contact geometry, when supplemented by additional physical properties, can be applied consistently throughout candidate generation and partition evaluation.

Here, we introduce VoroTIDE, a criterion-based protein-domain parser designed to address these requirements. VoroTIDE formulates Voronoi contact geometry into a pairwise *self-containment score F* to evaluate the coupling between structural fragments, and a global partition quality score G, *which is derived from F, to evaluate candidate partitions*. With VoroTIDE, we investigated whether protein-domain organization can be predicted and evaluated consistently using Voronoi-based criteria. We also compared VoroTIDE with widely used deep-learning and criterion-based parsers to assess both its predictive power and its limitations.

## Results

### Overview of the VoroTIDE framework

The central assumption of VoroTIDE is that, given an informative scoring function, there exists at least one partition along either the recursive splitting trajectory of the top-down search or the recursive merging trajectory of the bottom-up search that approximates the reference partition in terms of both the number of domains and their boundaries. This assumption motivates the implementation of many hierarchical searching parsers, including top-down recursive decomposition in PDP and PUU (Holm and Sander 1994, Alexandrov and Shindyalov 2003); bottom-up assembly in SWORD (Postic *et al*. 2017); protein units generated by Protein Peeling (Gelly, Brevern de, and Hazout 2006); and top-down splitting followed by bottom-up merging in UniDoc (Zhu *et al*. 2023).

However, these implementations have several limitations. First, top-down splitting and bottom-up merging are not generally inverse operations. Even when they use the same pairwise criterion and produce the same number of domains, they may yield different boundaries. Using only one trajectory can therefore exclude partitions that are accessible through the other. Second, when top-down and bottom-up procedures are applied sequentially, the second stage is constrained by the partition produced by the first. For example, a merge-only bottom-up stage cannot introduce a boundary that was missed during the preceding top-down stage. Third, the stopping criteria applied to each trajectory stop at the first partition that violates the criterion, although a later partition along the trajectory may satisfy it again.

VoroTIDE therefore performs independent bottom-up and top-down searches, exploring candidate partitions in opposite directions (**Figure 1**). The bottom-up process starts with individual structural units and recursively merges the contacting fragment pair with the highest self-containment score, *F*, until the whole structure forms a single fragment. The complementary top-down process starts from the complete structure and recursively applies the weakest admissible single or double cut with the lowest F until no further admissible division remains. In this study, we defined an admissible division as one that splits the current domain into smaller domains, each containing at least 40 residues. Every intermediate partition generated by both trajectories is retained as a candidate. After partition selection, fragments shorter than 40 residues or containing fewer than two regular SSEs are merged with the adjacent fragment with the highest *F*.

**Figure 1.**
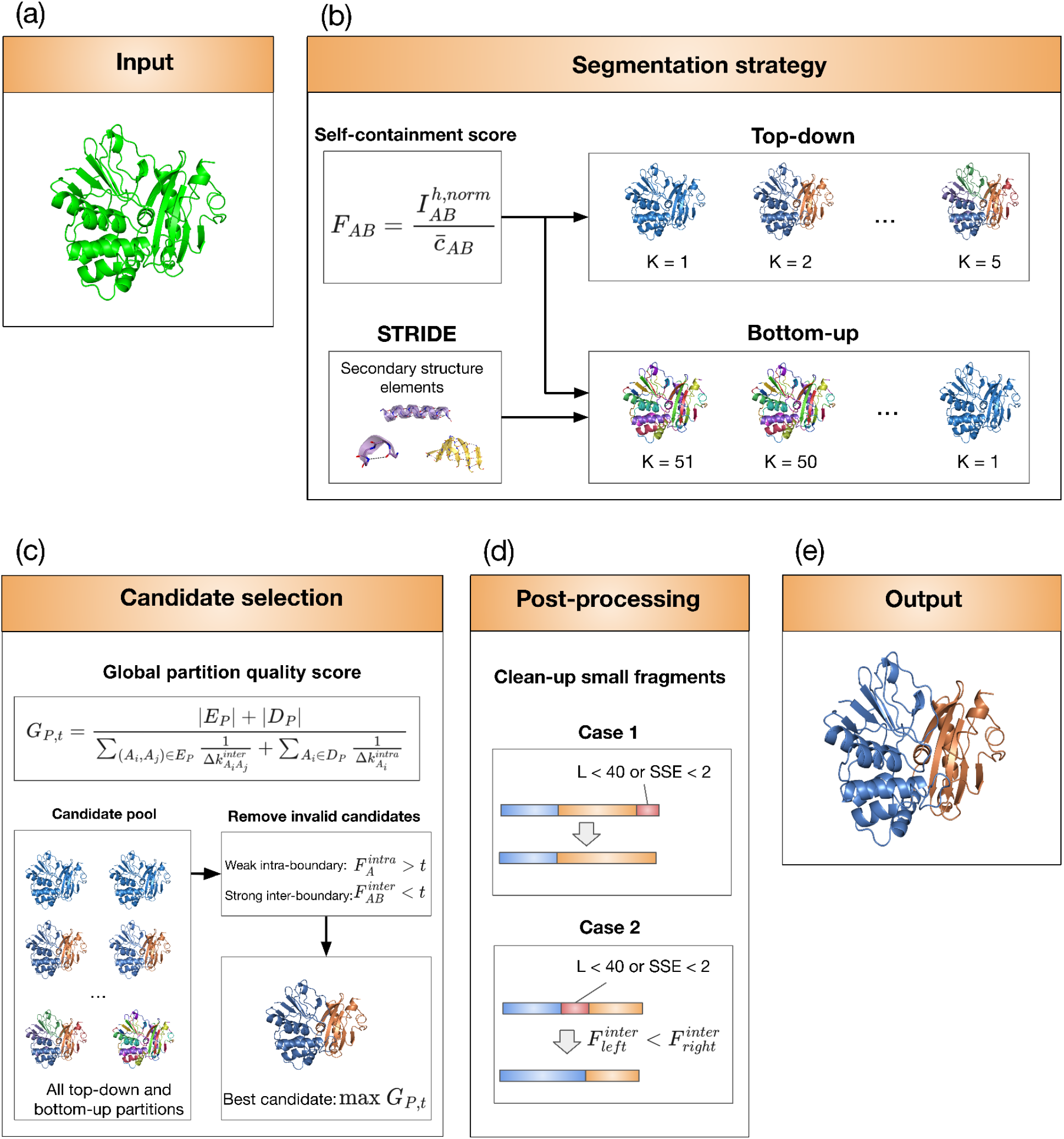
Overview of the VoroTIDE protein-domain segmentation pipeline. **(a)** The input protein 3D structure. **(b)** The 3D structure is represented using structural units derived from STRIDE assignments and Voronoi-derived structural contacts, then VoroTIDE explores the partition space through two independent hierarchical trajectories using the same pairwise self-containment score *F*. The top-down search recursively divides the structure at the most weakly coupled interfaces, whereas the bottom-up search starts from individual structural units and recursively merges the most strongly coupled fragments. All intermediate partitions generated by both trajectories are retained as candidates. **(c)** Candidate partitions are evaluated using the global partition-quality score *G*_*P,t*_, which simultaneously favors weak coupling between predicted domains and strong internal cohesion within each domain relative to the threshold *t*. Partitions that fail either condition are discarded, and the valid candidate with the highest *G*_*P,t*_ score is selected. **(d)** During post-processing, fragments shorter than 40 residues or containing fewer than two regular SSEs are merged with an adjacent fragment; when two neighboring fragments are available, the merge with the stronger pairwise coupling is selected. **(e)** The resulting partition constitutes the final VoroTIDE domain assignment.

Because both searches retain many candidates, VoroTIDE uses a global partition-quality score *G* derived from *F*. A good partition has low *F* between predicted domains and high *F* for the weakest admissible split within each domain. Both conditions are evaluated relative to threshold *t* (see Materials and Methods for details).

### Overview of the dataset construction

We constructed one tuning set and two benchmark sets with controls for training-set overlap, sequence redundancy, and structural redundancy (**Figure 2**). These controls are important for deep-learning parsers such as Chainsaw and Merizo, whose performance can be inflated by their related training structures (Lau, Kandathil, and Jones 2023, Wells *et al*. 2024).

**Figure 2.**
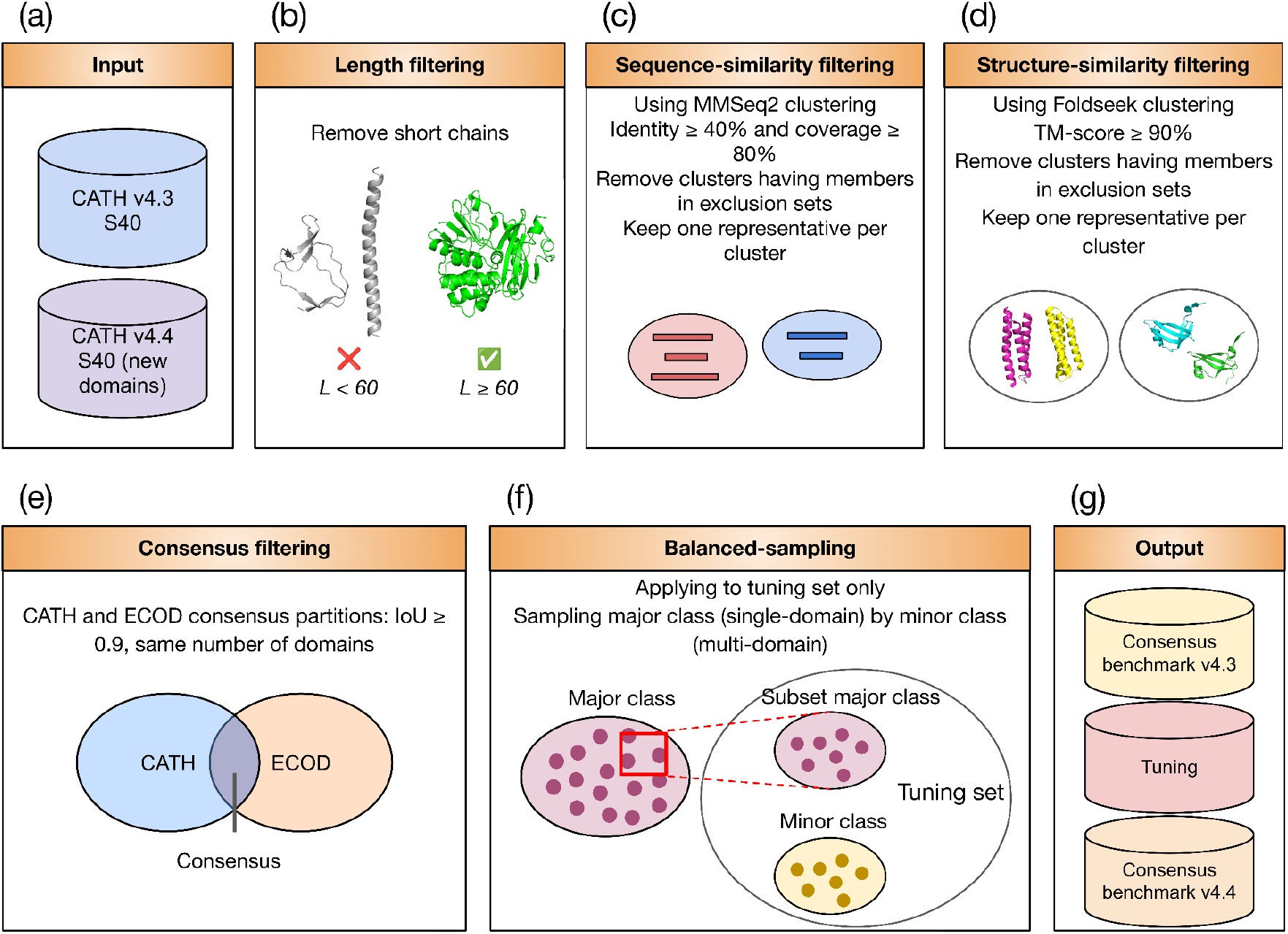
Dataset construction. **(a)** CONSENSUS-43 and tuning were derived from CATH v4.3 S40, whereas CONSENSUS-44 used domains newly represented in CATH v4.4. **(b)** The length filtering process removes structures having assigned-CATH region fewer than 60 residues. **(c)** After length filtering, candidate and exclusion chains were clustered with MMseqs2, and clusters containing exclusions and residual sequence hits were removed. **(d)** Foldseek was used to cluster the remaining structures to remove near-duplicates and those close to structures in the exclusions. **(e)** Chains were retained only when CATH and ECOD agreed on domain count and the partition score *IoU* was at least 0.90. **(f)** The tuning set was balanced between single- and multi-domain structures by downsampling the majority class (single-domain structures). **(g)** Finally, the three datasets CONSENSUS-43, tuning and CONSENSUS-44 were obtained.

CONSENSUS-43 was built from CATH v4.3 S40 (Sillitoe *et al*. 2020) after removing sequence and structural near-duplicates of the Chainsaw and Merizo training sets. We retained structures for which CATH and ECOD v295.2 (Schaeffer *et al*. 2026) agreed on the domain count and had a partition IoU ≥ 0.90, followed by final deduplication (see Materials and Methods for details).

The tuning set used the same CATH v4.3 pipeline, with CONSENSUS-43 as the exclusion set. After filtering, all multi-domain structures and an equal number of single-domain structures were retained to select the operating threshold *t*.

CONSENSUS-44 used the same pipeline on domains newly represented in CATH v4.4, excluding the Chainsaw and Merizo training sets, CONSENSUS-43, and the tuning set. Because the CATH v4.4 expansion of PDB domain annotations used Chainsaw (Waman et al. 2024), this benchmark is not fully independent of that method. CONSENSUS-44 was primarily used to assess the stability of tool performance rather than to rank evaluated domain parsers.

### VoroTIDE performance on the tuning dataset

The tuning set was used to select the threshold *t* that optimized both exact-*K* accuracy, which measures whether the predicted domain count is correct, and *K-score*, which reflects the magnitude of domain-count errors (see Materials and Methods). As the threshold *t* increased, VoroTIDE showed a clear trade-off between the two classes: single-domain performance decreased, whereas multi-domain performance improved before reaching a plateau (**Figure 3a, 3b**). Performance on the tuning set was broadly stable for *t* between 0.36 and 0.37, evaluated in increments of 0.005. Therefore, we adopted *t* = 0.365 within this plateau as the final operating threshold. At this value, the mean *K-score* was 0.950 and balanced exact-*K* accuracy was 83.4%, corresponding to 94.6% for single-domain and 72.3% for multi-domain structures.

**Figure 3.**
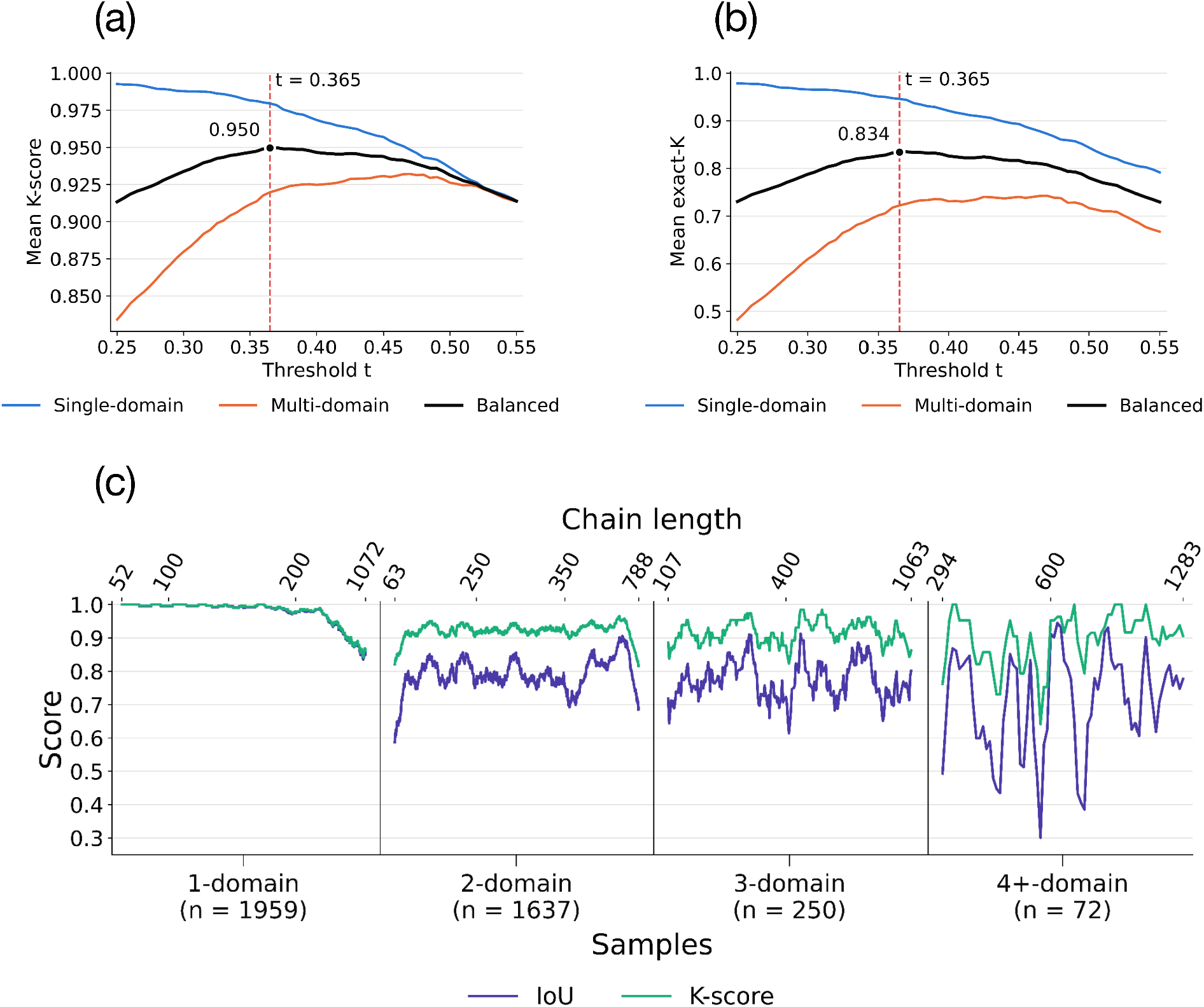
Threshold selection and performance of VoroTIDE on the tuning dataset. Relationship between **(a)** *mean K-score* or **(b)** *exact-K accuracy* and the threshold *t*. Blue and orange curves show the results for single-domain and multi-domain structures, respectively. The black curve represents the balanced performance, calculated as the arithmetic mean of the single-domain and multi-domain results. The vertical red dashed line indicates the selected threshold, *t = 0*.*365*. **(c)** Per-structure *IoU* (blue) and *K-score* (green) obtained at the selected threshold, grouped according to the target number of domains (1, 2, 3, and 4+ domains). Within each group, chains are ordered by chain length. The number of chains in each group is indicated below the corresponding category.

Additionally, VoroTIDE performance decreased as the domain count increased (**Figure 3c**). Average *exact-K* accuracy fell from 74.8% for two-domain structures to 64.0% for three-domain structures and 43.1% for structures with four or more domains, whereas the corresponding average *K-scores* remained comparatively high (0.921, 0.917 and 0.896). These results indicate that, although VoroTIDE increasingly failed to capture the exact domain count as domain organization became more complex, the magnitude of the counting errors generally remained limited. Detailed results for VoroTIDE on the tuning set are provided in Table S1 in Supplementary File 1.

### VoroTIDE performance on benchmark datasets

#### CONSENSUS-43

On the primary CONSENSUS-43 benchmark, VoroTIDE showed a clear class-dependent performance profile (**Figure 4**). For single-domain structures, VoroTIDE achieved the highest performance among all evaluated methods for both domain-count and partition-quality metrics, with an exact-*K* accuracy of 91.8% and a *K*-score of 0.968. Pairwise comparisons showed that these improvements over the other methods were statistically significant. For multi-domain structures, however, Chainsaw and Merizo achieved higher exact-*K* accuracies of 84.0% and 76.0%, respectively, compared with 71.0% for VoroTIDE, although the difference between VoroTIDE and Merizo was not statistically significant.

**Figure 4.**
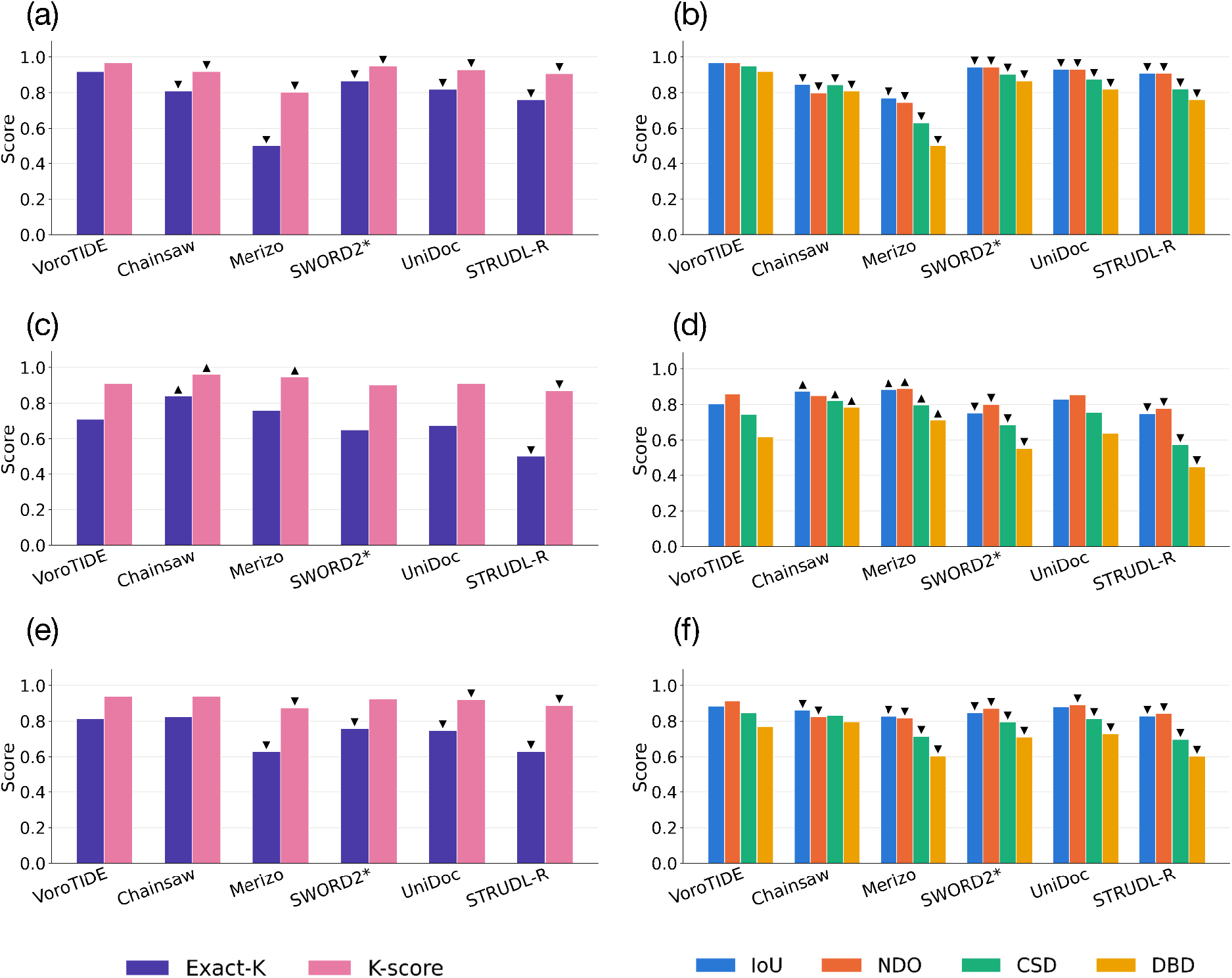
Performance of the evaluated domain parsers on CONSENSUS-43. Domain-count performance, measured by exact-*K* accuracy and K-score, is shown for **(a)** single-domain chains, **(c)** multi-domain structures, and **(e)** balanced performance across the two groups. Partition-quality performance, measured by *IoU, NDO, CSD*, and *DBD*, is shown for **(b)** single-domain chains, **(d)** multi-domain structures, and **(f)** balanced performance across the two groups. Balanced values were calculated as the arithmetic mean of the corresponding single-domain and multi-domain scores, giving equal weight to the two groups irrespective of their sample sizes. Triangles indicate pairwise differences from VoroTIDE that remained significant after Benjamini–Hochberg correction of paired-bootstrap *p-values* (q < 0.05). Upward triangles denote significantly better performance than VoroTIDE, whereas downward triangles denote significantly worse performance.

Among the criterion-based parsers, VoroTIDE remained competitive on multi-domain structures. Its domain-count performance was higher than that of SWORD2* and UniDoc, whereas STRUDL-R performed significantly worse. VoroTIDE also significantly outperformed SWORD2* and STRUDL-R on partition-quality metrics and was not significantly different from UniDoc. Specifically, VoroTIDE obtained *IoU, NDO, CSD*, and *DBD* scores of 0.804, 0.860, 0.744, and 0.618, respectively, compared with 0.829, 0.854, 0.755, and 0.640 for UniDoc. Thus, UniDoc was numerically higher for *IoU, CSD*, and *DBD*, whereas VoroTIDE was slightly higher for *NDO* and achieved better domain-count prediction. However, none of the partition-quality differences between the two methods was statistically significant.

Because CONSENSUS-43 is dominated by single-domain structures, we additionally evaluated balanced performance by assigning equal weight to the single- and multi-domain subsets (**Figure 4e, 4f**). Under this evaluation, VoroTIDE and Chainsaw were not significantly different for domain-count prediction, with balanced *exact-K accuracies* of 81.4% and 82.5% and *K-scores* of 0.939 and 0.940, respectively. VoroTIDE achieved the highest balanced *IoU* (0.885) and *NDO* (0.913), both significantly higher than Chainsaw, whereas the differences in *CSD* and *DBD* were not significant. Together, these results identified VoroTIDE and Chainsaw as the two strongest-performing methods under balanced evaluation. Detailed results for selected parsers on the CONSENSUS-43 set are provided in Tables S2, S3 and S4 in Supplementary File 1.

#### CONSENSUS-44

CONSENSUS-44 was used primarily to assess cross-release stability because CATH v4.4 is not fully independent of several evaluated parsers. VoroTIDE showed a performance profile similar to that observed on CONSENSUS-43: single-domain exact-*K* accuracy changed from 91.8% to 90.2%, multidomain accuracy from 71.0% to 74.0%, and balanced accuracy from 81.4% to 82.1% (**Figure 5**). Despite the slight decrease, VoroTIDE retained the highest single-domain performance among all evaluated methods. Among the criterion-based methods, VoroTIDE again achieved the strongest balanced performance and remained competitive with UniDoc on multidomain structures. Specifically, for domain-count prediction, VoroTIDE achieved an exact-*K* accuracy of 74.0% and a *K*-score of 0.925, compared with 69.3% and 0.911 for UniDoc, respectively. For partition quality, VoroTIDE obtained IoU, NDO, CSD, and DBD scores of 0.834, 0.879, 0.794, and 0.623, respectively, whereas UniDoc obtained 0.831, 0.869, 0.759, and 0.640. Thus, VoroTIDE was numerically higher on domain-count prediction and on three of the four partition-quality metrics, whereas UniDoc achieved a slightly higher DBD score. None of these differences reached statistical significance.

**Figure 5.**
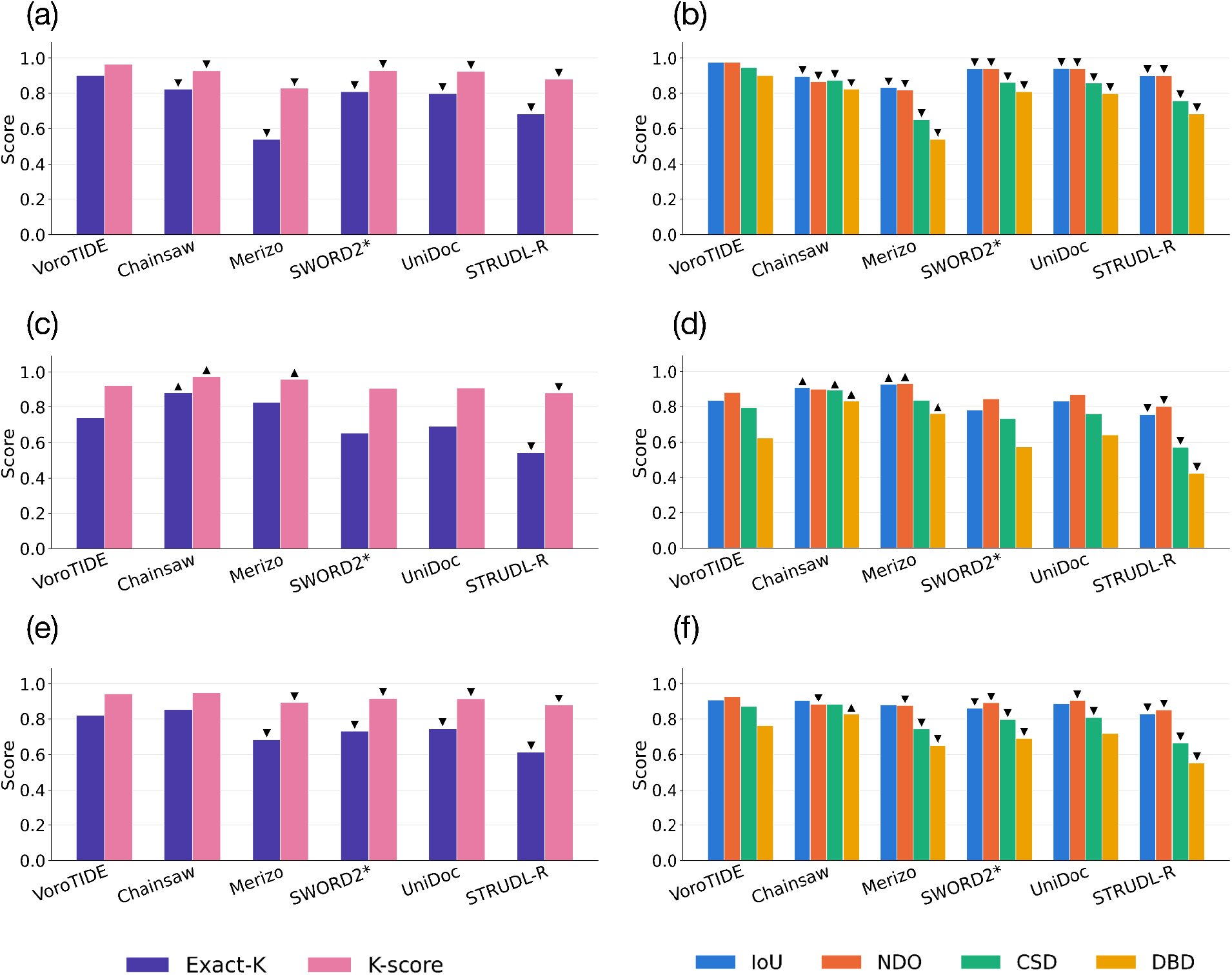
Performance of the evaluated domain parsers on CONSENSUS-44. Domain-count performance, measured by exact-*K accuracy* and *K-score*, is shown for **(a)** single-domain chains, **(c)** multi-domain structures, and **(e)** balanced performance across the two groups. Partition-quality performance, measured by *IoU, NDO, CSD*, and *DBD*, is shown for **(b)** single-domain chains, **(d)** multi-domain structures, and **(f)** balanced performance across the two groups. Balanced values were calculated as the arithmetic mean of the corresponding single-domain and multi-domain scores, giving equal weight to the two groups irrespective of their sample sizes. Triangles indicate pairwise differences from VoroTIDE that remained significant after Benjamini–Hochberg correction of paired-bootstrap *p-values* (q < 0.05). Upward triangles denote significantly better performance than VoroTIDE, whereas downward triangles denote significantly worse performance.

When considering multidomain structures alone, the deep-learning parsers Chainsaw and Merizo outperformed VoroTIDE. However, when single- and multi-domain performance were weighted equally, VoroTIDE outperformed Merizo and remained competitive with Chainsaw. Thus, the main performance patterns were consistent across the two benchmarks. Detailed results for selected parsers on the CONSENSUS-44 set are provided in Tables S5, S6 and S7 in Supplementary File 1.

#### Representative case studies

We also examined representative cases to compare VoroTIDE’s behavior with two other widely used parsers: Chainsaw, a deep-learning method, and UniDoc, a criterion-based method. We selected examples spanning four agreement patterns with the CATH reference (**Figure 6**): cases in which all three parsers were correct, cases in which only VoroTIDE was correct, cases in which only VoroTIDE was incorrect, and cases in which none reproduced the reference partition. Single- and multi-domain structures were included for each pattern, providing a qualitative view of both VoroTIDE-specific successes and failure modes. A predicted partition was considered correct when it had the correct number of domains and a partition IoU ≥ 0.90. A prediction was considered incorrect if it had the wrong number of domains. Details of reference domain assignments and predictions from all evaluated parsers for each benchmark sample are provided in Supplementary File 2.

**Figure 6.**
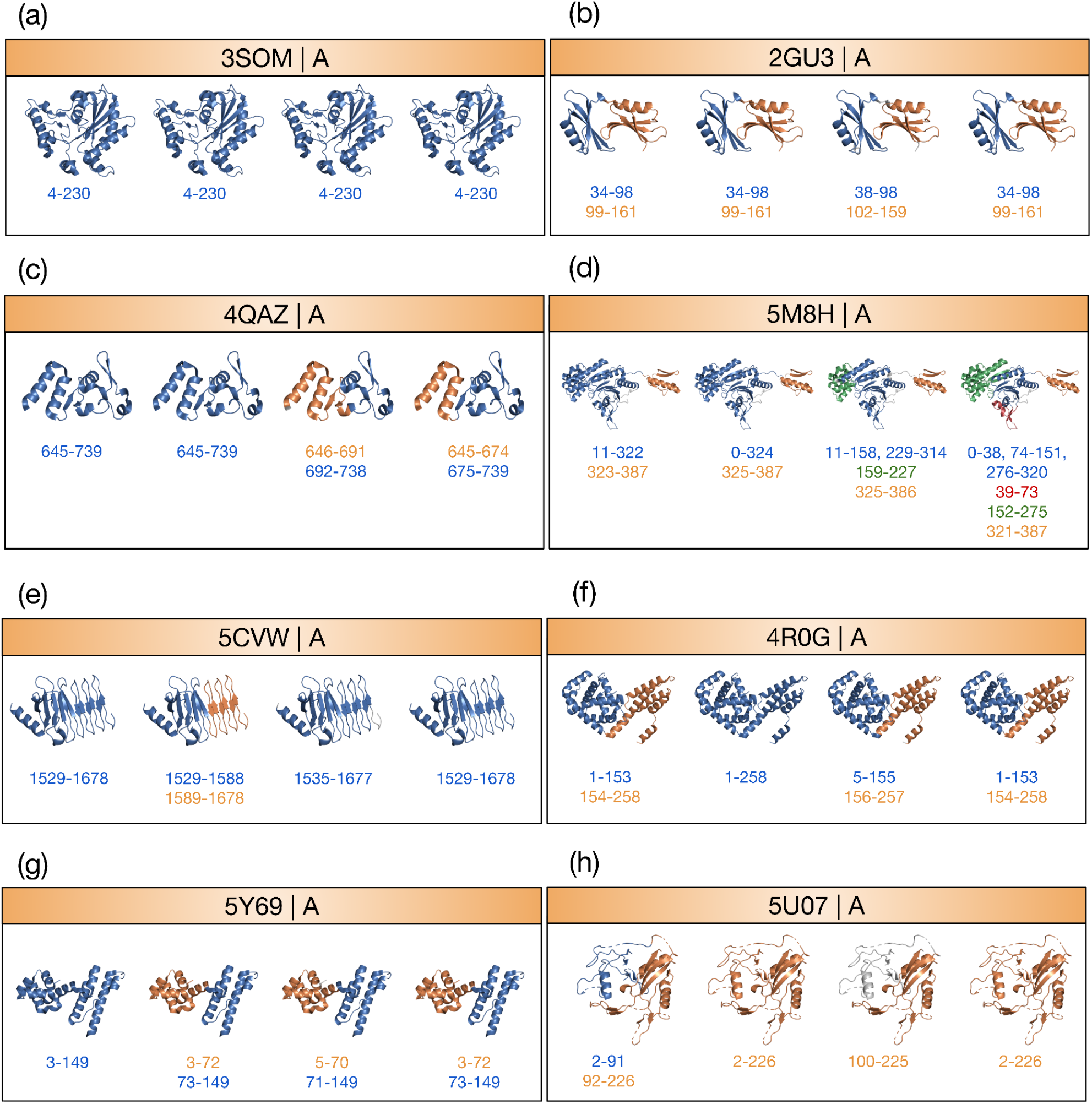
Representative domain partitions. In each panel, CATH, VoroTIDE, Chainsaw, and UniDoc are shown from left to right, respectively. Panels show cases where all parsers are correct (**a, b**), only VoroTIDE is correct (**c, d**), only VoroTIDE is incorrect (**e, f**), or none reproduces the reference (**g, h**), with single- and multi-domain structures for each pattern. Colors identify corresponding domains. Residues absent from the PDB were bridged only for display. Residues omitted by a parser are those that have been unassigned by that parser.

In cases where all parsers were correct, 3SOM chain A is a compact single α/β domain that all three methods kept intact (**Figure 6a**), whereas 2GU3 chain A contains two similarly sized domains separated by a short loop that was recovered by all methods (**Figure 6b**). VoroTIDE and UniDoc reproduced the reference boundary exactly, while Chainsaw placed the cut within the same loop but left nine linker residues unassigned.

In cases where only VoroTIDE was correct, the errors of the comparison methods were mainly due to over-segmentation. The small single-domain protein 4QAZ chain A was preserved intact by VoroTIDE, whereas Chainsaw split the chain near its center and UniDoc separated an N-terminal α-hairpin (**Figure 6c**). In the two-domain protein 5M8H chain A, all methods detected the principal interdomain boundary, but Chainsaw and UniDoc further subdivided the large N-terminal domain, whereas VoroTIDE retained the correct two-domain organization (**Figure 6d**).

Conversely, VoroTIDE failed in examples where a structurally distinct region was either too weakly integrated or too strongly coupled. In 5CVW chain A, VoroTIDE incorrectly separated an N-terminal β-rich segment from an otherwise single-domain structure, whereas Chainsaw and UniDoc kept the chain intact (**Figure 6e**). In 4R0G chain A, VoroTIDE missed the reference boundary between two helical domains and merged the chain into one domain, while UniDoc and Chainsaw recovered the reference separation (**Figure 6f**).

Finally, some structures were inconsistent with all three parsers. In 5Y69 chain A, all methods divided the CATH single-domain reference into two helical bundles at nearly the same position, suggesting a reproducible structural subdivision not represented in the reference annotation (**Figure 6g**). This case is particularly ambiguous because with visual inspection, these two helical bundles are spatially separated. In 5U07 chain A, all methods failed to recover the two-domain reference: VoroTIDE and UniDoc merged the entire chain, whereas Chainsaw detected only the C-terminal domain and left the irregular N-terminal region unassigned (**Figure 6h**).

## Discussion

VoroTIDE shows that much of the curated domain organization can be recovered from an explicit physical self-containment criterion without learning from domain annotations, especially for single-domain structures. On CONSENSUS-43, it had the strongest balanced performance among the criterion-based methods and domain-count performance comparable to Chainsaw. Its strong single-domain results are consistent with the method: a chain remains unsplit when no admissible internal cut is weak enough. The lower accuracy at higher domain counts indicates that recovering the annotated domain organization becomes more challenging, especially if the domains are tightly packed.

Voronoi geometry and *F* serve different roles. Voronoi adjacency defines direct local contacts, while *F* asks whether an interface is weak relative to fragment size and internal packing. This normalization is needed because the raw contact area is not comparable across fragments of different sizes. Thus, Voronoi defines the interactions and *F* turns them into a domain-level decision variable.

Multi-domain structures were more challenging not only for VoroTIDE but also for other explicit structure-based parsers, whereas deep-learning parsers like Chainsaw and Merizo showed a clearer advantage. Physical criteria assume that annotated boundaries correspond to a predefined form of structural decoupling. However, curated assignments can also reflect higher-order context, evolution, homology, or classification conventions (Veretnik *et al*. 2004, Schaeffer *et al*. 2021). Deep-learning parsers trained on these annotations may therefore implicitly capture some of these additional patterns (Lau, Kandathil, and Jones 2023, Wells *et al*. 2024). There might still be physical signals, but they may not be limited to the explicit contact and physicochemical terms used for VoroTIDE or other selected criterion-based parsers in this study.

The multi-domain gap also shows VoroTIDE’s complementary role. Each boundary of VoroTIDE is linked to an explicit interface, its hydropathy-weighted coupling, and the internal cohesion of adjacent fragments. Disagreements with curated annotations are therefore directly interpretable via physical criteria. The similar profile on CONSENSUS-44 suggests that this behavior is stable across CATH releases, although that set is not fully independent of several evaluated parsers. Overall, physical self-containment accounts for a substantial portion of curated domain organization, while the remaining multi-domain gap points to information not captured by the present interface criterion.

Since the formula of *F* comprises multiple components, we conducted an ablation study on the tuning set to assess the contributions of its individual components by comparing alternative variants (see “Ablation study” section of Supplementary File 1). For each variant, the threshold *t* was retuned on the tuning set. Evaluation on the CONSENSUS-43 set showed that most components of *F*, including hydropathy and size-dependent terms, contributed to the performance. Removing any of these components led to a slight decrease in balanced domain-count performance, although the differences were not significant. These results suggest that much of the domain signal is captured by the Voronoi contact area, while other physical criteria provide supplementary information.

The only exception was variant 3, in which the exponent β was set to 1. We therefore further examined the effect of β by testing a range of values from 0.1 to 2.0 on the tuning set (**Figure S1** in Supplementary File 1). VoroTIDE’s domain-number prediction performance remained on a plateau between β = 0.7 and 1.6, a range that includes both 0.86 and 1. When β fell below 0.7, performance on multi-domain structures declined rapidly, resulting in a sharp drop in balanced performance. Above β = 1.6, performance on both single- and multi-domain structures gradually declined, resulting in lower balanced performance. A possible explanation for this trend is how the effective threshold *t* scales with fragment size. Specifically, when β is small, the threshold barely depends on size. Since the contact between large fragments is usually large and that between small fragments is usually small, large domains are over-merged, while small domains are over-split. In contrast, when β is large, the threshold grows faster with size than the interfaces. The contact between large fragments of the same domain then falls below the expected area of their size, thus large domains become over-split. Small fragments easily exceed their much lower expected area, leading to small domains being over-merged.

## Materials and Methods

### Structural representation

VoroTIDE describes a protein chain in terms of elementary structural units and geometric contacts. The structural units are contiguous helical, strand, and coil segments derived from secondary-structure assignments computed using STRIDE (Frishman and Argos 1995). Each unit is treated as indivisible during both hierarchical searches. These units initialize the bottom-up hierarchy and constrain the admissible cuts in the top-down search.

Geometric contacts are represented by an all-heavy-atom, additively weighted Voronoi tessellation of atomic balls computed with Voronota v1.29 using its atomic-radius model and a 1.4 Å probe (Olechnovič and Venclovas 2014). Only heavy atoms from standard ATOM records are included in the tessellation; HETATM records, including modified residues, are excluded from the contact graph and mapped back only when residue ranges are reconstructed for output. Two atoms are direct geometric neighbors when their Voronoi cells share a facet, and the facet area quantifies the geometric extent of their contact. Atomic facet areas are summed to residue–residue contact areas and then reused throughout the search: interfaces between structural units or larger fragments are obtained by summing the corresponding residue–residue areas, so the tessellation is performed only once per chain.

### Pairwise self-containment score

A structural boundary between two domains *A* and *B* is expected to correspond to relatively weak geometric coupling, reflected by a limited Voronoi contact interface 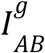 (**Figure 7d**). We further hypothesize that such weak interfaces can also be less hydrophobically stabilized. Specifically, a limited contact area may provide less opportunity to bury nonpolar residues from the solvent, and contacts across a domain boundary are expected, on average, to be more polar than contacts contributing to the interior of a compact structural domain (**Figure 7e**).

**Figure 7.**
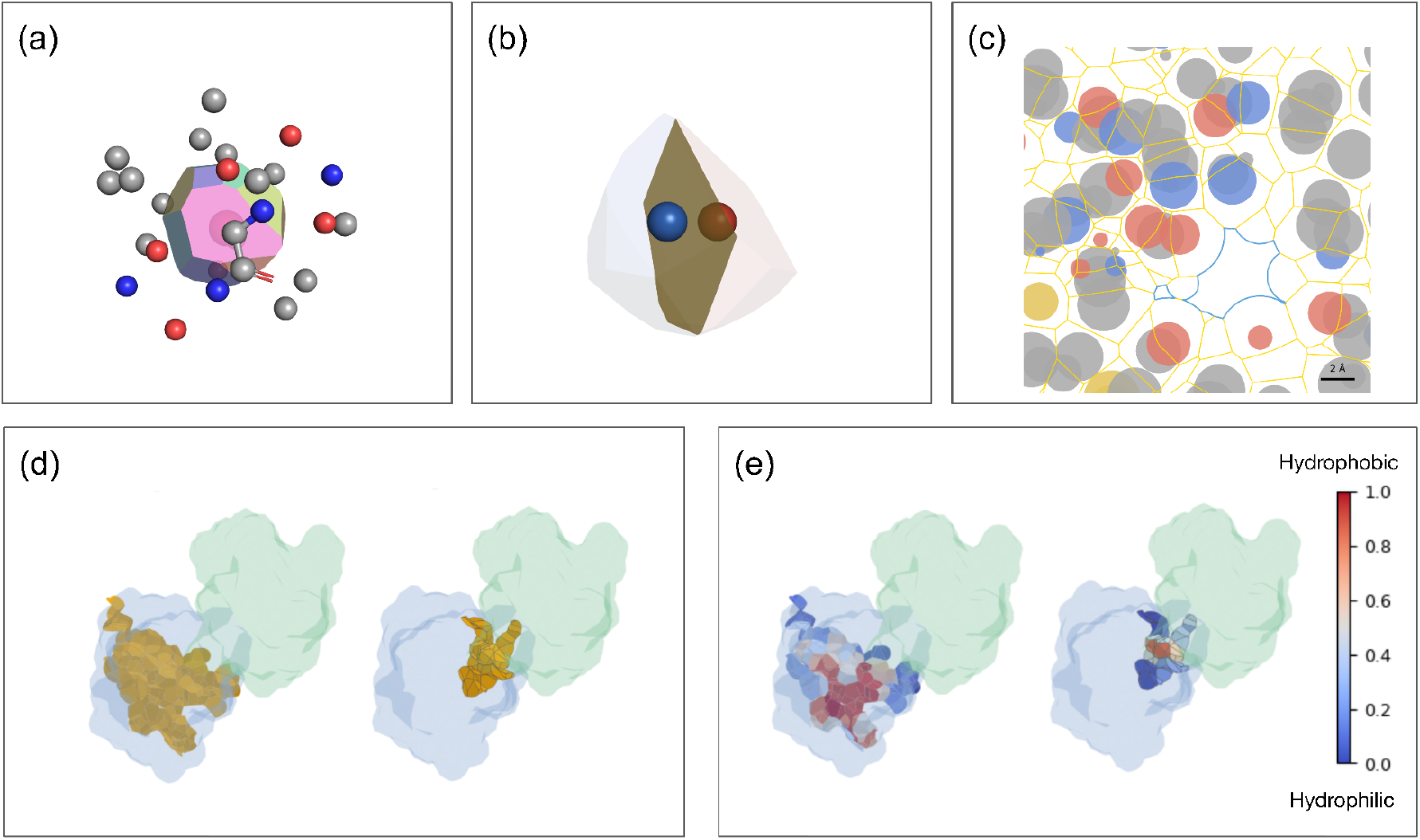
Voronoi contact geometry and hydropathy weighting. **(a)** Atomic Voronoi cell. **(b)** Shared facet between neighboring atoms; its area measures geometric contact. **(c)** Contact facets in a multi-atom system. **(d)** Geometric contact interfaces for an intra-domain cut (left) and an inter-domain boundary (right). Yellow surfaces highlight the Voronoi contact facets between the two fragments, shown as translucent blue and green regions. **(e)** The corresponding interfaces are colored by normalized hydropathy, ranging from blue (0, most hydrophilic) through white to red (1, most hydrophobic).

VoroTIDE therefore weights each residue–residue Voronoi contact by residue hydropathy. We use the Kyte–Doolittle scale (Kyte and Doolittle 1982), normalized from its original range [−4.5, 4.5] to [0,1]:

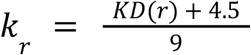

For two contacting residues *r* and *s* sharing a Voronoi contact area 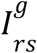, the contact area is weighted by multiplying with their mean normalized hydropathy. The hydropathy-weighted interface 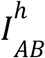 between fragments *A* and *B* is therefore calculated as:

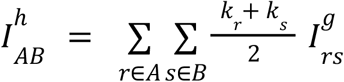

Equivalently, the hydropathy-weighted interface 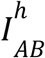 can be interpreted as the geometric Voronoi interface 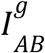 multiplied by an effective hydropathy factor *k*_*AB*_, representing the contact-area-weighted mean hydropathy of the interacting residues between two fragments A and B:

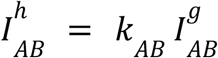

Consequently, 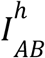 depends on both the geometric interface 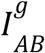 and the effective hydropathy factor *k*_*AB*_. When both 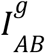 and *k*_*AB*_ are small, which means the contact interface is limited and formed predominantly by hydrophilic residues, 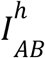 is small, consistent with our assumptions of a domain boundary. Conversely, an increase in any of these factors will increase 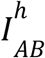 and therefore weaken the evidence that the interface between *A* and *B* corresponds to a domain boundary.

The absolute value of 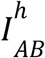, however, depends on the sizes of the interacting fragments. For example, bigger fragments can have naturally larger contact areas, while smaller fragments tend to have smaller contact areas. To account for this effect, we introduce a reduced-size term for two fragments A and B, containing *N*_*A*_ and *N*_*B*_ residues respectively, defined as:

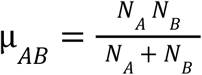

The reduced-size term µ corresponds to the harmonic mean of *N*_*A*_ and *N*_*B*_ with the constant factor of two omitted, as this constant only rescales the normalization without affecting its relative behavior. When the two fragments differ substantially in size, *N*_*A*_ >> *N*_*B*_ for example, µ is governed primarily by the smaller fragment, *B* with *N*_*B*_, whereas for fragments of similar size, both contribute comparably. This property prevents the size of a very large fragment from artificially reducing the normalized interface score when it interacts strongly with a much smaller fragment, as could occur with arithmetic- or geometric-mean-based normalizations. The hydropathy-weighted interface 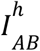 is then normalized by an effective reduced-size term:

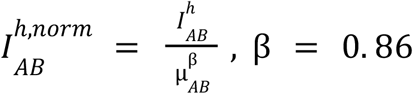

The exponent β controls how the reduced-size term µ_*AB*_ scales the contact interface with fragment size. In PDP (Alexandrov and Shindyalov 2003), the corresponding normalization is 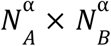, with α = 0. 43. For fragments of comparable size: *N*_*A*_ ≈ *N*_*B*_ ≈ *N*, the reduced-size term satisfies µ_*AB*_≈ *N*/2, so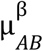 scales as *N*^β^ up to a constant factor, whereas the PDP normalization scales as *N* ^2α^. We therefore set β = 2α = 0. 86, preserving the same size-scaling behavior in the comparable-size limit.

The normalized interface 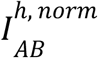 also needs to be evaluated relative to the overall structural packing of fragments *A* and *B*. We therefore define the geometric contact-based normalization term:

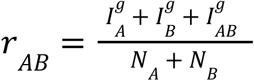

Here, 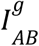 is the total geometric contact area between residues belonging to fragments A and B. For each fragment *X* ∈ {*A,B*}, 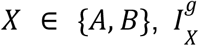 is calculated by summing geometric contact areas between distinct elementary structural units within *X*, counting each contact once. Contacts within individual units are excluded from this sum. If the inter-unit sum is zero, including when *X* consists of a single unit, the sum of within-unit geometric contact areas is used instead. This convention is applied consistently in both hierarchical searches and in the later evaluation of candidate partitions.

Finally, we define the pairwise self-containment score *F*_*AB*_ which serves as the scoring function in VoroTIDE for evaluating the coupling strength between two structural fragments:

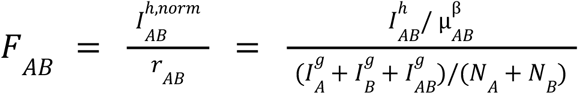

A high value of *F*_*AB*_ indicates that fragments *A* and *B* are strongly coupled relative to their internal structural reference, providing evidence that they should remain within the same domain. Conversely, a low *F*_*AB*_ indicates a weak interface relative to the internal packing of the fragments and therefore supports the presence of a domain boundary between them.

We also created different variants of *F* by removing specific components to evaluate their performance. Details of each variant are provided in the “Ablation study” section of Supplementary File 1.

### Global partition quality score

VoroTIDE ranks the retained candidates with a global criterion derived from F that combines inter-domain separation and intra-domain cohesion.

For a candidate partition *P* = {*A*_1_, *A*_2_, …, *A*_*k*_}, we assume that the coupling between any two distinct domains *A*_*i*_ and *A*_*j*_ should be weaker than the coupling associated with any admissible intra-domain cut within either domain. Let *C*(*A*_*i*_) denote the set of admissible bipartitions (*A*_*i*,1_, *A*_*i*,2_) of *A*_*i*_, where each resulting fragment contains at least 40 residues. If *C*(*A*_*i*_) is non-empty, the weakest admissible intra-domain coupling of *A* is then defined as:

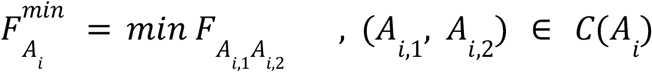

Internal bipartitions use the same single- and double-cut configurations as those used in top-down search. Domains shorter than 80 residues have no admissible split and contribute no intra-domain margin.

Then according to the assumption, we have:

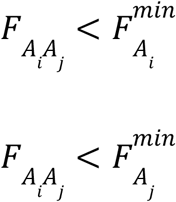

Where 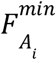 and 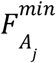 denote the weakest admissible intra-domain coupling values of *A*_*i*_ and *A*_*j*_, respectively. Therefore, we seek a common threshold *t* that distinguishes inter-domain interfaces from internal domain cuts. Such a threshold, if it exists, must satisfy:

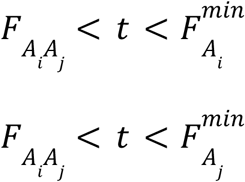

We then define the inter-domain and intra-domain scores as:

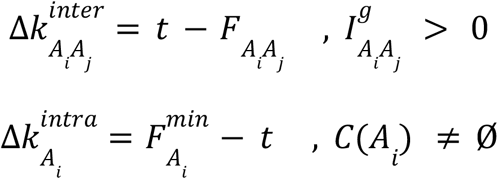

The inter-domain score is calculated only between two domains *A*_*i*_ and *A*_*j*_ that are in contact, i.e., 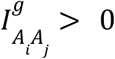, whereas the intra-domain score is calculated for every domain *A*_*i*_ in P for which *C*(*A*_*i*_) is non-empty. For compact notation, we define *E*_*p*_ as the set of contacting domain pairs and *D* as the set of domains with at least one admissible intra-domain cut:

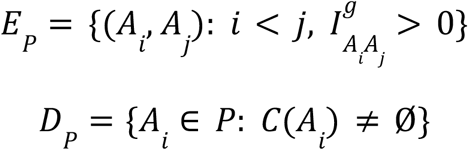

Finally, the global partition quality score for a candidate partition *P* at threshold *t* is defined as the harmonic mean of the inter-domain and intra-domain scores:

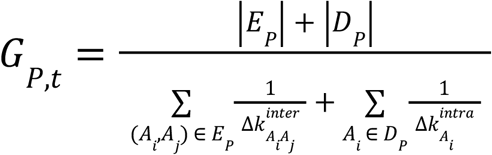

We use the harmonic mean to limit the extent to which large margins can compensate for small margins elsewhere in the partition. Consequently, *G*_*P,t*_ is strongly penalized when any included margin is small and favors partitions in which the inter-domain and intra-domain scores are consistently high. If any inter-domain or intra-domain score is non-positive, the partition *P* is considered invalid and is discarded before the calculation of the global partition quality score. If the structure contains fewer than 80 residues or none of the candidate partitions satisfy the validity criterion, VoroTIDE assigns the entire structure as a single domain.

### Independent top-down and bottom-up searches

VoroTIDE generates candidate partitions through two independent hierarchical searches operating on the same indivisible structural units, including helical, strand, and coil segments. The bottom-up search starts with each unit as a separate fragment and progressively merges contacting fragments. The top-down search starts from the entire chain and progressively splits fragments at boundaries between units, subject to the minimum-size constraints.

The top-down search starts from the entire chain and recursively divides fragments at their weakest admissible interfaces. For each fragment, its constituent units are ordered along the sequence, and two types of bipartition are evaluated. A single cut separates this ordered list into two groups, each containing at least 40 residues. A double cut separates an internal group from the union of the two flanking groups, allowing the identification of a domain inserted within another domain. In this configuration, the internal group and each flanking group must individually contain at least 40 residues. Among admissible cuts with a nonzero geometric interface, the cut with the lowest *F* is retained for each fragment. At each step, the fragment whose retained cut has the lowest *F* is divided, generating a new partition. This process continues until no fragment has an admissible cut.

The bottom-up search starts with each elementary structural unit as a separate fragment. At each step, the contacting pair with the highest *F* is merged, and the contact areas and sizes associated with the resulting fragment are updated. Merging is not restricted to sequence-adjacent fragments, allowing domains composed of discontinuous sequence segments to form. The process continues until a single fragment remains or no contacting pair is available for merging.

The initial partition and every partition produced along each trajectory are retained in a candidate pool. The two trajectories can yield different boundaries even with the same number of fragments, thereby providing complementary candidate partitions. These candidates are subsequently evaluated using the global partition quality score G, with threshold t, as described above.

### Post-processing

Post-processing is applied only after candidate selection and comprises two successive steps. First, sequence segments shorter than 40 residues are reassigned to a sequence-adjacent domain, starting with the shortest segment and updating the segmentation after each reassignment. Second, domains containing fewer than two regular SSEs (helices or strands) are merged into a sequence-adjacent domain, starting with the smallest domain. In both steps, the recipient domain is selected by the highest pairwise score *F* among contacting sequence-adjacent candidates. If none of these candidates is in contact, the nearest candidate along the sequence is used. Processing stops when no eligible segment or domain remains, or only one domain is left.

### Evaluation metrics

To evaluate the performance of protein domain parsers, we considered two complementary aspects: the accuracy of the predicted domain count and the accuracy of domain localization. Different metrics were used to evaluate each aspect.

### Domain-count evaluation

To assess the accuracy of the predicted number of domains, we first considered the *exact K accuracy* metric. It has a score of 1 when the predicted number of domains *K*_*pred*_ exactly matches that of the target partition *K*_*target*_, and 0 otherwise.

However, *exact-K* accuracy is the proportion of chains for which the predicted domain count matches the reference count. Each partition contributes 1 for a correct count and 0 otherwise. This binary contribution does not reflect the magnitude of a counting error. Therefore, we introduced the *K-score*, which is based on Bray–Curtis dissimilarity, to quantify how far the predicted domain count is from the target (Bray and Curtis 1957):

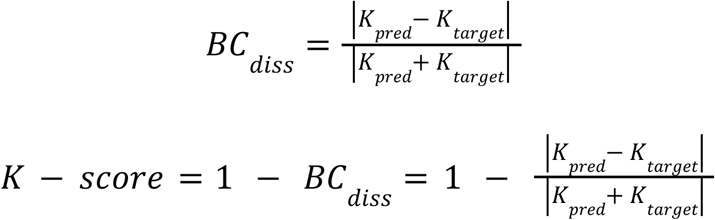

Where *BC*_*diss*_ is the Bray–Curtis dissimilarity between the number of target and predicted domains. The *K-score* ranges from 0 to 1. A value of 1 is obtained when the predicted and target partitions contain the same number of domains, whereas a value of 0 is obtained when one partition contains at least one assigned domain while the other contains no assigned domain, i.e., all residues are treated as linker or unassigned.

The *K-score* equals 1 for the correct count and decreases as the relative counting error increases. For example, with a three-domain target, predicting four domains yields 0.857, whereas predicting ten yields 0.462.

### Partition-quality evaluation

For partition-quality evaluation, we used four metrics: normalized domain overlap (NDO; Tai *et al*. 2005), intersection over union (IoU; Lau, Kandathil, and Jones 2023), chain segment distance (CSD; Le *et al*. 2025), and domain boundary distance (DBD; Tress *et al*. 2007). In principle, NDO and IoU quantify the agreement between target and predicted partitions based on residue overlap between corresponding domains, whereas CSD and DBD are distance-based metrics that evaluate their positional agreement at the chain-segment and domain-boundary levels, respectively.

### Statistical analysis

Pairwise differences between VoroTIDE and each comparison parser were evaluated on matched chains using a two-sided paired bootstrap with 3000 resamples. For balanced analyses, each bootstrap replicate sampled equal numbers of single- and multi-domain structures with replacement within each stratum before averaging the two class-specific means. Bootstrap *p-values* were adjusted over the full pairwise comparison table using the Benjamini–Hochberg procedure, with *q* < 0.05 considered significant. *Exact-K* conclusions were additionally checked with exact McNemar tests, and continuous metrics with Wilcoxon signed-rank tests on the non-balanced subsets; their *p-values* were also Benjamini–Hochberg-adjusted within the corresponding sensitivity-test family. These served as sensitivity analyses, while the bootstrap remained the primary test because the reported summary statistics are means.

### Evaluated domain parsers

VoroTIDE was compared with UniDoc (Zhu *et al*. 2023), SWORD2 (Cretin *et al*. 2022), Chainsaw (Wells *et al*. 2024), and Merizo (Lau, Kandathil, and Jones 2023). UniDoc and SWORD2 are criterion-based parsers. Chainsaw and Merizo are supervised deep-learning methods. All tools were obtained from their official repositories. For SWORD2, we used only the top-ranked “optimal partition” from its candidate pool for each structure, denoted as SWORD2*. Predictions were mapped to the same reference residues before scoring. Empty outputs or execution failures were scored as empty predictions rather than excluded.

In addition, we reconstructed STRUDL, which also used weighted Voronoi contact areas defined by radical planes and Kernighan–Lin graph partitioning (Wernisch, Hunting, and Wodak 1999). Since no public code or active server was found, we denoted our reconstructed version of it as STRUDL-R in this study. The logistic classifier in STRUDL-R was fitted on the tuning set using predefined structural descriptors.

### Data preprocessing

We constructed three datasets sequentially: CONSENSUS-43, tuning, and CONSENSUS-44. CONSENSUS-43 and tuning were derived from chains for which all constituent domains were represented in the CATH v4.3 S40 non-redundant set (Sillitoe et al. 2020). CONSENSUS-44 was instead derived from chains whose constituent domains were newly released in the CATH v4.4 S40 set (Waman *et al*. 2024) and thus were absent from the CATH v4.3 (Sillitoe *et al*. 2020). The tuning dataset was used to select the VoroTIDE threshold *t*, while CONSENSUS-43 served as the primary benchmark for comparing the performance of the evaluated domain parsers. Because CATH v4.4 incorporates Chainsaw into its PDB domain-segmentation workflow and uses Chainsaw, Merizo, and UniDoc in its broader automated segmentation framework (Waman *et al*. 2024), annotations in this release are not fully independent of the evaluated deep-learning parsers. CONSENSUS-44 was therefore used as a secondary benchmark to assess whether VoroTIDE’s performance on CONSENSUS-43 remained stable across two CATH releases, rather than ranking the methods.

All three datasets followed the same filtering logic, differing only in their respective exclusion sets. They were constructed through four successive stages: 1. Length filtering; 2. Sequence-similarity filtering and removal of clusters associated with the corresponding exclusion set; 3. Structure-similarity filtering and removal of clusters associated with the exclusion set; 4. CATH–ECOD consensus filtering. The tuning dataset included an additional balanced-sampling stage, in which equal numbers of single-domain and multi-domain structures were selected.

#### Length filtering

Candidate structures were required to have a CATH-assigned region of at least 60 residues. This left 35581 candidates for CONSENSUS-43/tuning and 17278 for CONSENSUS-44.

#### Sequence-similarity filtering

Candidates and exclusions were clustered together with MMseqs2 at 40% identity and 80% coverage (Steinegger and Söding 2017). Clusters containing an exclusion chain were removed, and one representative was kept from each remaining cluster. Exclusions were the Chainsaw/Merizo training sets for CONSENSUS-43, the CONSENSUS-43 benchmark set for tuning, and all preceding sets plus the training sets for CONSENSUS-44. This left 4350, 16332, and 1755 candidates, respectively.

#### Structure-similarity filtering

After removing structures without ECOD annotation, Foldseek was used to remove structural near-duplicates (Kempen van *et al*. 2024). A match was excluded at TM-score ≥ 0.90 normalized by the longer chain, or at sequence identity ≥ 0.90 with TM-score ≥ 0.80. The same rule retained one representative within each set. This left 4051, 15555, and 1487 candidates, respectively.

#### CATH–ECOD consensus filtering

Finally, we applied consensus filtering between CATH and ECOD using the fixed ECOD v295.2 release (Schaeffer *et al*. 2026). A protein chain was retained only when the two classifications assigned the same number of domains and their partitions had an *IoU* of at least 0.90 on its CATH-assigned region. A final residual de-duplication pass was then applied to the retained chains. The resulting CONSENSUS-43 benchmark contained 3231 structures, including 3031 single-domain and 200 multi-domain structures, whereas CONSENSUS-44 contained 1060 structures, including 933 single-domain and 127 multi-domain structures. For the tuning dataset, consensus filtering produced 11241 candidates, comprising 9282 single-domain and 1959 multi-domain structures, before balanced sampling.

#### Balanced-sampling

Since the tuning dataset was imbalanced toward single-domain structures, we applied balanced sampling to reduce potential bias toward the majority class when defining the operating range of *t*. Specifically, we randomly sampled 1959 single-domain structures to match the 1959 multi-domain structures available in the candidate pool. The resulting tuning dataset therefore contained 3918 structures with equal proportions of single-domain and multi-domain structures.

## Abbreviations and symbols

VoroTIDE: Voronoi-based Two-way Independent Domain Exploration
CATH: Class–Architecture–Topology–Homologous superfamily classification
ECOD: Evolutionary Classification of Protein Domains
SSE: secondary-structure element
NDO: normalized domain overlap
IoU: intersection over union
DBD: domain-boundary distance
CSD: chain segment distance
F: VoroTIDE pairwise self-containment score
G: Global partition quality score
t: partition-selection threshold
μ: reduced-size term

## Supplementary material description

**Supplementary File 1** contains average performance results for VoroTIDE on the tuning set and for all evaluated parsers on CONSENSUS-43 and CONSENSUS-44, together with the ablation study.

**Supplementary File 2** is an Excel workbook containing per-chain reference domain boundaries, predicted domain assignments, and evaluation metrics for the parsers evaluated on each dataset.

## Data availability

Benchmark manifests, per-chain domain assignments, metric tables and scripts used to generate the reported results are deposited at GitHub. CATH and ECOD annotations remain subject to their respective database terms and should be obtained from the original resources.

## Code availability and accessibility statement

VoroTIDE source code, installation instructions and command-line examples are available on GitHub. The software is intended to be freely accessible to academic users. The repository included the exact release corresponding to the manuscript, dependency information and scripts for reproducing the benchmark analyses.

## Acknowledgments

The authors thank Andrey Kajava for discussions on Voronoi- and SASA-based structural analysis that helped motivate the development of the scoring function. The author also acknowledges the computational resources and institutional support provided by CRBM, CNRS, and the University of Montpellier.

## Author contributions

Quoc Khang Le: Software; Writing – review and editing; Validation; Methodology; Conceptualization; Formal analysis; Data curation; Visualization. Guillaume Postic: Review; Conceptualization; Validation; Supervision. Eric Angel: Review; Supervision. Fariza Tahi: Review; Supervision.

