## Supplementary File 1 for "VoroTIDE: Protein domain segmentation through independent hierarchical searches using Voronoi tessellation and physical self-containment"

### Supplementary Tables

**Table S1:** Performance of VoroTIDE with selected threshold $t$ = 0.365 on all structures of the tuning set.

| Class | N | IoU | NDO | CSD | DBD | Exact-K (%) | K-score |
|---|---|---|---|---|---|---|---|
| Single | 1959 | 0.978 | 0.978 | 0.965 | 0.946 | 94.6 | 0.980 |
| Multi | 1959 | 0.785 | 0.849 | 0.753 | 0.566 | 72.3 | 0.920 |
| Balanced | 3918 | 0.882 | 0.914 | 0.859 | 0.756 | 83.4 | 0.950 |
| 2-domain | 1637 | 0.790 | 0.855 | 0.763 | 0.561 | 74.8 | 0.921 |
| 3-domain | 250 | 0.772 | 0.831 | 0.717 | 0.605 | 64.0 | 0.917 |
| 4+-domain | 72 | 0.707 | 0.780 | 0.653 | 0.546 | 43.1 | 0.896 |

**Table S2:** Performance of the evaluated domain parsers on single-domain structures of the CONSENSUS-43 set.

| Tool | IoU | NDO | CSD | DBD | Exact-K (%) | K-score |
|---|---|---|---|---|---|---|
| VoroTIDE | **0.967** | **0.967** | **0.949** | **0.918** | **91.8** | **0.968** |
| Chainsaw | 0.846 | 0.799 | 0.844 | 0.809 | 80.9 | 0.919 |
| Merizo | 0.770 | 0.745 | 0.631 | 0.499 | 49.9 | 0.803 |
| SWORD2* | 0.943 | 0.943 | 0.904 | 0.867 | 86.7 | 0.948 |
| UniDoc | 0.931 | 0.930 | 0.875 | 0.819 | 81.9 | 0.929 |
| STRUDL-R | 0.908 | 0.908 | 0.821 | 0.760 | 76.0 | 0.908 |

**Table S3:** Performance of the evaluated domain parsers on multi-domain structures of the CONSENSUS-43 set.

| Tool | IoU | NDO | CSD | DBD | Exact-K (%) | K-score |
|---|---|---|---|---|---|---|
| VoroTIDE | 0.804 | 0.860 | 0.744 | 0.618 | 71.0 | 0.910 |
| Chainsaw | 0.875 | 0.850 | **0.820** | **0.784** | **84.0** | **0.962** |
| Merizo | **0.883** | **0.890** | 0.798 | 0.712 | 76.0 | 0.946 |
| SWORD2* | 0.752 | 0.800 | 0.686 | 0.554 | 65.0 | 0.902 |
| UniDoc | 0.829 | 0.854 | 0.755 | 0.640 | 67.5 | 0.910 |
| STRUDL-R | 0.748 | 0.778 | 0.575 | 0.447 | 50.0 | 0.869 |

**Table S4:** Performance of the evaluated domain parsers on all structures of the CONSENSUS-43 set, with balanced performance calculated across single- and multi-domain structures.

| Tool | IoU | NDO | CSD | DBD | Exact-K (%) | K-score |
|---|---|---|---|---|---|---|
| VoroTIDE | **0.885** | **0.913** | **0.847** | 0.768 | 81.4 | 0.939 |
| Chainsaw | 0.861 | 0.825 | 0.832 | **0.797** | **82.5** | **0.940** |
| Merizo | 0.827 | 0.817 | 0.715 | 0.606 | 62.9 | 0.875 |
| SWORD2* | 0.848 | 0.872 | 0.795 | 0.710 | 75.8 | 0.925 |
| UniDoc | 0.880 | 0.892 | 0.815 | 0.729 | 74.7 | 0.920 |
| STRUDL-R | 0.828 | 0.843 | 0.698 | 0.603 | 63.0 | 0.888 |

**Table S5:** Performance of the evaluated domain parsers on single-domain structures of the CONSENSUS-44 set.

| Tool | IoU | NDO | CSD | DBD | Exact-K (%) | K-score |
|---|---|---|---|---|---|---|
| VoroTIDE | **0.978** | **0.978** | **0.947** | **0.902** | **90.2** | **0.966** |
| Chainsaw | 0.897 | 0.865 | 0.874 | 0.822 | 82.2 | 0.929 |
| Merizo | 0.830 | 0.818 | 0.651 | 0.540 | 54.0 | 0.828 |
| SWORD2* | 0.940 | 0.940 | 0.860 | 0.808 | 80.8 | 0.931 |
| UniDoc | 0.941 | 0.940 | 0.857 | 0.797 | 79.7 | 0.925 |
| STRUDL-R | 0.901 | 0.901 | 0.757 | 0.683 | 68.3 | 0.881 |

**Table S6:** Performance of the evaluated domain parsers on multi-domain structures of the CONSENSUS-44 set.

| Tool | IoU | NDO | CSD | DBD | Exact-K (%) | K-score |
|---|---|---|---|---|---|---|
| VoroTIDE | 0.834 | 0.879 | 0.794 | 0.623 | 74.0 | 0.925 |
| Chainsaw | 0.910 | 0.902 | **0.893** | **0.832** | **88.2** | **0.975** |
| Merizo | **0.929** | **0.934** | 0.835 | 0.761 | 82.7 | 0.960 |
| SWORD2* | 0.781 | 0.843 | 0.733 | 0.574 | 65.4 | 0.908 |
| UniDoc | 0.831 | 0.869 | 0.759 | 0.640 | 69.3 | 0.911 |
| STRUDL-R | 0.755 | 0.800 | 0.571 | 0.424 | 54.3 | 0.879 |

**Table S7:** Performance of the evaluated domain parsers on all structures of the CONSENSUS-44 set, with balanced performance calculated across single- and multi-domain structures.

| Tool | IoU | NDO | CSD | DBD | Exact-K (%) | K-score |
|---|---|---|---|---|---|---|
| VoroTIDE | **0.906** | **0.928** | 0.871 | 0.763 | 82.1 | 0.946 |
| Chainsaw | 0.904 | 0.883 | **0.883** | **0.827** | **85.2** | **0.952** |
| Merizo | 0.879 | 0.876 | 0.743 | 0.651 | 68.3 | 0.894 |
| SWORD2* | 0.860 | 0.892 | 0.796 | 0.691 | 73.1 | 0.919 |
| UniDoc | 0.886 | 0.905 | 0.808 | 0.719 | 74.5 | 0.918 |
| STRUDL-R | 0.828 | 0.851 | 0.664 | 0.554 | 61.3 | 0.880 |

### Ablation study

To evaluate the contribution of different components in F, we conducted an ablation study. Specifically, we created several variants of F by removing some of its components and evaluated their performance on the CONSENSUS-43 dataset.

Variant 1: remove hydropathy effect:

$$F_{AB}^{Variant\ 1} = \frac{I_{AB}^{g,\ norm}}{r_{AB}} = \frac{I_{AB}^{g} / \mu_{AB}^{\beta}}{(I_A^g + I_B^g + I_{AB}^g)/(N_A + N_B)}$$

Variant 2: remove geometric contact-based normalization term:

$$F_{AB}^{Variant\ 2} = I_{AB}^{h,\ norm} = I_{AB}^{h} / \mu_{AB}^{\beta}$$

Variant 3: remove exponent β (set β = 1):

$$F_{AB}^{Variant\ 3} = \frac{I_{AB}^{h} / \mu_{AB}}{r_{AB}} = \frac{I_{AB}^{h} / \mu_{AB}}{(I_A^g + I_B^g + I_{AB}^g)/(N_A + N_B)}$$

Variant 4: remove size-dependent terms $\mu_{AB}^{\beta}$ and $(N_A + N_B)$:

$$F_{AB}^{Variant\ 4} = \frac{I_{AB}^{h}}{(I_A^g + I_B^g + I_{AB}^g)}$$

Variant 5: remove size-dependent terms $\mu^{\beta}_{AB}$, $(N_A + N_B)$ and hydropathy effect:

$$F^{Variant\ 5}_{AB} = \frac{I^g_{AB}}{(I^g_A + I^g_B + I^g_{AB})}$$

**Table S8:** Performance of the selected *F* formulation and its variants, each evaluated using a threshold *t* selected on the tuning set, on single-domain structures in CONSENSUS-43.

| Variant | t | IoU | NDO | CSD | DBD | Exact-K (%) | K-score |
|---|---|---|---|---|---|---|---|
| Selected F | 0.365 | **0.967** | **0.967** | **0.949** | **0.918** | **91.8** | **0.968** |
| Variant 1 | 0.822 | 0.954 | 0.954 | 0.932 | 0.894 | 89.4 | 0.957 |
| Variant 2 | 16.110 | 0.954 | 0.954 | 0.929 | 0.888 | 88.8 | 0.955 |
| Variant 3 | 0.214 | 0.961 | 0.961 | 0.938 | 0.902 | 90.2 | 0.962 |
| Variant 4 | 0.051 | 0.958 | 0.958 | 0.919 | 0.878 | 87.8 | 0.952 |
| Variant 5 | 0.099 | 0.957 | 0.957 | 0.915 | 0.868 | 86.8 | 0.949 |

**Table S9:** Performance of the selected *F* formulation and its variants, each evaluated using a threshold *t* selected on the tuning set, on multi-domain structures in CONSENSUS-43.

| Variant | t | IoU | NDO | CSD | DBD | Exact-K (%) | K-score |
|---|---|---|---|---|---|---|---|
| Selected F | 0.365 | 0.804 | 0.86 | 0.744 | 0.618 | 71.0 | 0.910 |
| Variant 1 | 0.822 | 0.82 | 0.854 | 0.743 | 0.628 | 70.0 | 0.914 |
| Variant 2 | 16.110 | **0.845** | **0.866** | **0.758** | **0.630** | 71.5 | **0.922** |
| Variant 3 | 0.214 | 0.82 | 0.859 | 0.741 | 0.622 | **72.5** | 0.917 |
| Variant 4 | 0.051 | 0.751 | 0.819 | 0.678 | 0.515 | 59.0 | 0.880 |
| Variant 5 | 0.099 | 0.739 | 0.821 | 0.681 | 0.508 | 57.0 | 0.870 |

**Table S10:** Performance of the selected *F* formulation and its variants, each evaluated using a threshold *t* selected on the tuning set, on all structures in CONSENSUS-43, with balanced performance calculated across single- and multi-domain structures.

| Variant | t | IoU | NDO | CSD | DBD | Exact-K (%) | K-score |
|---|---|---|---|---|---|---|---|
| Selected F | 0.365 | 0.885 | **0.913** | **0.847** | **0.768** | **81.4** | 0.939 |
| Variant 1 | 0.822 | 0.887 | 0.904 | 0.838 | 0.761 | 79.7 | 0.936 |
| Variant 2 | 16.110 | **0.9** | 0.91 | 0.843 | 0.759 | 80.2 | 0.938 |
| Variant 3 | 0.214 | 0.89 | 0.91 | 0.84 | 0.762 | **81.4** | **0.94** |
| Variant 4 | 0.051 | 0.854 | 0.889 | 0.798 | 0.697 | 73.4 | 0.916 |
| Variant 5 | 0.099 | 0.848 | 0.889 | 0.798 | 0.688 | 71.9 | 0.91 |

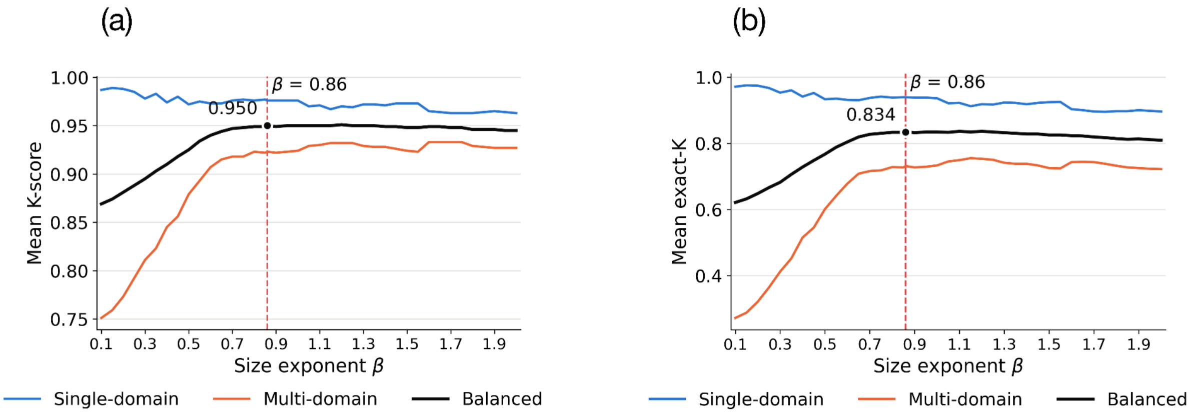


**Figure S1. Exponent selection and performance of VoroTIDE on the tuning dataset.** Relationship between **(a)** *mean K-score* or **(b)** *exact-K accuracy* and the exponent $\beta$. Blue and orange curves show the results for single-domain and multi-domain structures, respectively. The black curve represents the balanced performance, calculated as the arithmetic mean of the single-domain and multi-domain results. The vertical red dashed line indicates the selected exponent, $\beta = 0.86$.
